# Ligand-induced ubiquitination regulates endocytosis and homeostasis of the ERECTA receptor kinase for stomatal development

**DOI:** 10.1101/2025.06.29.662054

**Authors:** Liangliang Chen, Minh Huy Vu, Alicia M. Cochran, Crystal F. Ying, Keiko U. Torii

## Abstract

Stomata, valves on the plant epidermis, control gas and water vapor exchange. The patterning and spacing of stomata are primarily regulated by the ERECTA leucine-rich repeat receptor kinase (LRR-RK). Ubiquitination is a crucial mechanism to regulate the homeostasis of RKs by impacting their protein stability and localization. It has been shown that plant U-box ubiquitin E3 ligases, PUB30 and PUB31, act as key attenuators of the ERECTA signaling pathway. However, the molecular link between ubiquitination and function of ERECTA remains unclear. Here, we reveal that perception of the peptide ligand EPIDERMAL PATTERNING FACTOR2 (EPF2) by ERECTA induces the K63-linked ubiquitination of ERECTA mediated by PUB30 and PUB31. We further identify the specific ubiquitination sites within the juxtamembrane and kinase domains of ERECTA. Importantly, the site-directed mutagenesis of these K63-linked ubiquitination sites overly inhibited stomatal development, indicating that the ubiquitination-deficient ERECTA is hyperactive. The ubiquitination-deficient ERECTA stably accumulates at the plasma membrane and fails to internalize upon EPF2 application. Our findings thus uncover a mechanism by which ligand-induced ubiquitination orchestrates receptor homeostasis and endocytosis for proper stomatal patterning and differentiation.

## Introduction

Precise regulation of stomatal patterning optimizes plant gas exchange and water-use efficiency (Pillitteri & Torii, 2012). In *Arabidopsis thaliana*, this process is tightly controlled by cell-cell signaling mediated receptor-like kinases (RLKs, those with known ligands are referred hereafter as receptor kinases (RKs)). Among them, the ERECTA family plays a central role. ERECTA, together with its paralogs ERL1 and ERL2, negatively regulates stomatal formation by perceiving a family of secreted peptide ligands, EPIDERMAL PATTERNING FACTORs (EPFs) and EPF-LIKEs (Hara *et al*., 2007; Hara *et al*., 2009; Hunt & Gray, 2009; Kondo *et al*., 2010; Sugano *et al*., 2010). Among the ERECTA-family members, ERECTA is expressed in protodermal- and early stomatal-lineage cells. ERECTA primarily perceives EPF2 and activates a downstream mitogen-activated protein kinase (MAPK) signaling cascade, leading to the inhibition of the master transcription factors of stomatal development (Lampard *et al*., 2008; Lee *et al*., 2012; Horst *et al*., 2015). Consequently, loss-of-function mutations in *ERECTA* result in excessive asymmetric cell divisions, thereby increasing the number of stomatal precursor cells, meristemoids (Shpak *et al*., 2005; Lee *et al*., 2012). In addition to its role in stomatal patterning, ERECTA regulates aerial organ growth, notably inflorescence and pedicel elongation (Torii *et al*., 1996; Shpak *et al*., 2004; Shpak, 2013). Here, ERECTA expressed in the internal tissue perceives EPFL4/6 to promote cell proliferation in a non-cell autonomous manner (Uchida *et al*., 2012). While the downstream signaling pathway for ERECTA-mediated stem/pedicel elongation is less established, it also mediates the MAPK cascade (Meng *et al*., 2012).

Studies in the past decade revealed the mechanism of ERECTA receptor activation. ERECTA constitutively forms a receptor complex with TOO MANY MOUTHS (TMM) (Nadeau & Sack, 2002; Lee *et al*., 2012; Lee *et al*., 2015). Before ligand perception, the activity of ERECTA is inhibited by BRI1-KINASE INHIBITOR1 (BKI1) (Wang *et al*., 2017; Chen *et al*., 2025). Perception of EPF2 peptides recruits co-receptors BRI1-ASSOCITED KINASE (BAK1)/SOMATIC EMBRYOGENESIS RECPTOR KINASE1 (SERK1) and SERK2 (Lee *et al*., 2012; Meng *et al*., 2012; Lee *et al*., 2015; Meng *et al*., 2015; Lin *et al*., 2017). BAK1 subsequently transphosphorylates the C-terminal tail domain of ERECTA. This triggers eviction of BKI1 from ERECTA and recruitment of ubiquitin E3 ligases, PLANT U-BOX 30 (PUB30) and PUB31, which in turn downregulate the activated ERECTA (Chen *et al*., 2023; Chen *et al*., 2025). This intricate mechanism of ERECTA attenuation also operates for inflorescence/pedicel elongation. Consistently, loss-of-function *pub30pub31* double mutant plants exhibit greatly reduced number of stomata and extremely elongated inflorescence and pedicels due to excessive accumulation of ERECTA proteins (Chen *et al*., 2023).

Protein ubiquitination (also known as ubiquitylation) has been recognized as a central mechanism in regulating both abundance and subcellular localization of plasma membrane (PM) proteins (Leitner *et al*., 2012; Dubeaux *et al*., 2018; Romero-Barrios & Vert, 2018). Canonically, lysine 48 (K48)-linked polyubiquitination marks target proteins for proteasomal degradation, including degradation of some membrane proteins in mammals (Foot *et al*., 2017). In plants, K63-linked polyubiquitination has emerged as a key mechanism for membrane protein degradation through impacting their subcellular localizations (Romero-Barrios & Vert, 2018). For example, K63 polyubiquitination of the flagellin receptor FLS2 and the brassinosteroid hormone receptor BRI1 promotes their internalization and vacuolar degradation via multivesicular bodies (MVBs), providing an important means of fine-tuning the duration of receptor signaling (Lu *et al*., 2011; Zhou *et al*., 2018). As mentioned above, we have previously identified PUB30 and PUB31 as the E3 ubiquitin ligases that mediate the ubiquitination of ERECTA in response to peptide ligand stimulation (Chen *et al*., 2023; Chen *et al*., 2025). However, the exact sites or types of polyubiquitination as well as the mechanistic consequence of ERECTA ubiquitination on its intracellular trafficking remain largely unexplored.

In this study, we investigated the functional significance of ERECTA ubiquitination for its protein behaviors and phenotypic consequences. For this purpose, we surveyed publicly available ubiquitome data as well as inference from the studies of other plant RKs and generated ubiquitination-deficient ERECTA variants, in which three lysine residues (K617, K625, and K668) were substituted by arginine. We examined the effects of this mutant receptor on stomatal development, ubiquitination, intracellular trafficking of ERECTA, and the receptor protein abundance. Our results demonstrate that K63-linked ubiquitination of ERECTA is essential for its ligand-induced endocytosis and eventual vacuolar degradation. Mutagenesis of these three residues results in reduced endocytosis of ERECTA, increased receptor abundance, and a hyperactive inhibition of stomatal development. These findings uncover a critical role for ubiquitination in constraining ERECTA signaling output and provide mechanistic insight into how post-translational regulation governs receptor function during plant development.

## Results

### Ubiquitination-deficient ERECTA overly inhibits stomatal development

The loss-of-function *erecta* mutant exhibits increased number of small stomatal lineage cells (Shpak *et al*., 2005; Lee *et al*., 2012). In contrast, the *pub30pub31* double mutant overly reduces stomatal development (Chen *et al*., 2023). We hypothesized that the expression of ubiquitination-deficient ERECTA in the *erecta* null mutant (*er-105*) background leads to over-inhibition of stomatal development. A previous large-scale profiling of the Arabidopsis membrane-protein ubiquitome has identified a single lysine residue (K668) within the ERECTA kinase domain as a ubiquitination site (Grubb *et al*., 2021). With this in mind, we generated an epitope-tagged version of ERECTA in which K668 was substituted with arginine via site-directed mutagenesis (ERECTA_K668R)_ and conducted a functional complementation assay. However, *ERECTApro::ERECTA_K668R_-FLAG* failed to confer the over-complementation phenotype, indicating that there are other functionally relevant ubiquitination sites in ERECTA (Fig. S1).

To further explore potential sites, we performed sequence alignment between ERECTA and BRI1, the latter of which is known to be ubiquitinated at K866 in the juxtamembrane domain (Martins *et al*., 2015). Two additional lysine residues (K617, K625) in the juxtamembrane domain of ERECTA were identified (Fig. 1A, B). We subsequently mutated each site in combination with the previously reported site (K668) to generate higher-order ERECTA variants (ERECTA_K617RK668R_ or ERECTA_K625RK668R_). Both *ERECTApro::ERECTA_K617RK668R_-FLAG* and *ERECTApro::ERECTA_K625RK668R_-FLAG* produced reduced number of stomata and meristemoid compared to the control *ERECTApro:: ERECTA-FLAG* lines, suggesting that these two residues are additional ubiquitination sites in ERECTA (Fig. S1). We therefore substituted all three lysine residues with arginine residues (ERECTA_K617RK625RK668R_, hereafter ERECTA_3KR_). As expected, the expression of *ERECTApro::ERECTA_3KR_-FLAG* led to over-rescue phenotypes in stomatal development and pedicel elongation, with significantly reduced SMI (stomatal + meristemoid index = (number of stomata + meristemoid)/total number of epidermal cells x 100) and vastly increased pedicel length (Fig. 1C). Consistently, the expression of *ERECTApro::ERECTA_3KR_-YFP* also led to repressed stomatal development and enhanced pedicel growth (Fig. S2). Transcript levels of the transgenes are comparable among the transgenic lines either expressing *ERECTA - FLAG* and *ERECTA_3KR_-FLAG* (Fig. S3A) or *ERECTA -YFP* and *ERECTA_3KR_-YFP* (Fig. S3B), indicating that the observed excessive phenotypic rescues are not attributable to the transgene overexpression. Combined, the data suggests that multiple lysine residues cooperatively regulate ERECTA activity via ubiquitination and that ubiquitination on ERECTA plays a positive role in stomatal development.

**Figure 1.**
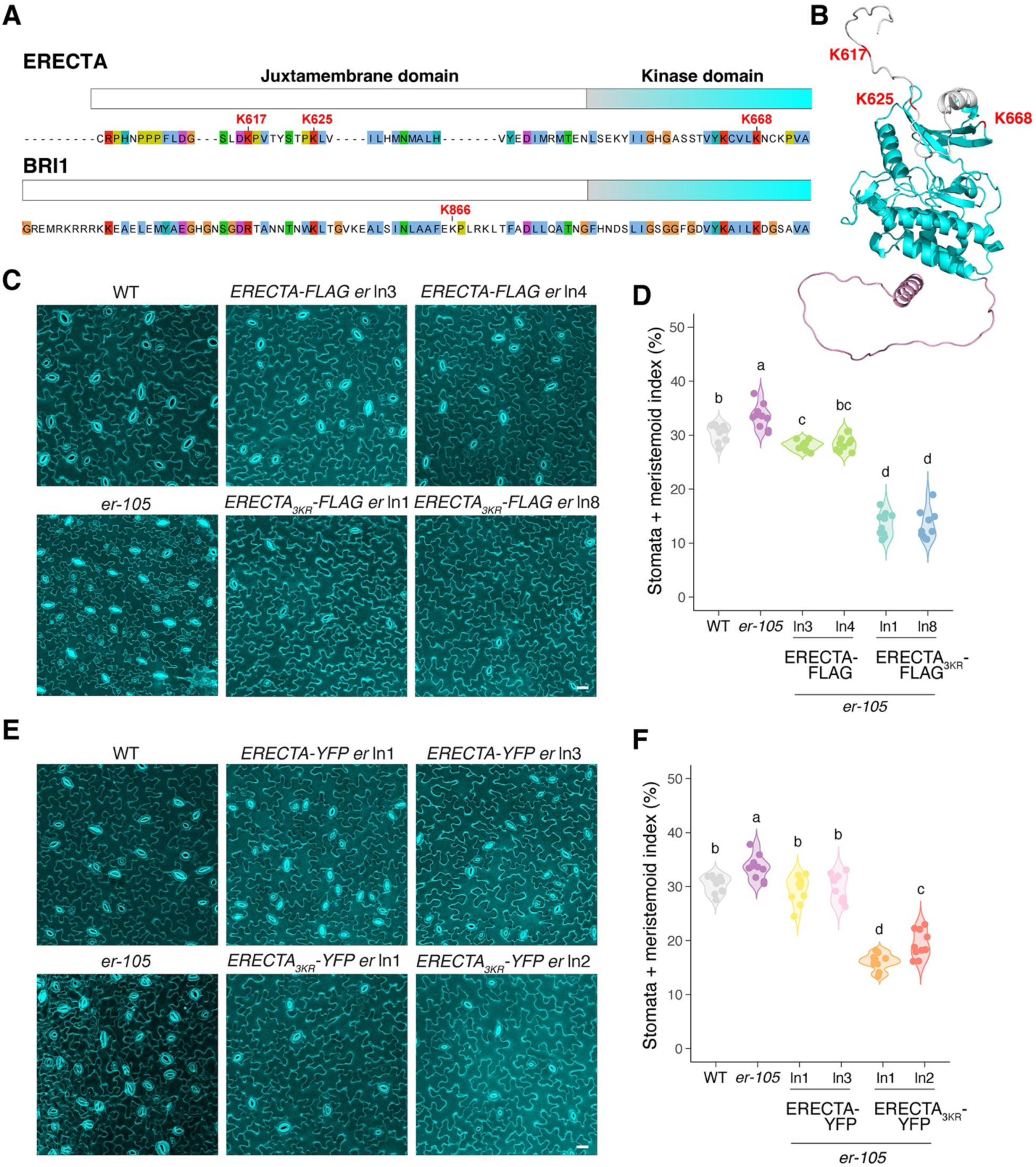
Ubiquitination-deficient ERECTA overly inhibits stomatal development. **(A)** Sequence alignment of the cytoplasmic domain of ERECTA (ERECTA_CD) and BRI1 (BRI1_CD). Juxtamembrane domains are highlighted in white, and the Kinase domains are in cyan. The selected lysine residues are labeled. **(B)** Structural modeling of the cytoplasmic domain of ERECTA, with the Juxtamembrane domain in light gray, and the Kinase domain in cyan. **(C)** Representative confocal microscopy of cotyledon abaxial epidermal from 7-day-old wild type (WT), *er-105* (*er*), *ERECTA-FLAG er*, and *ERECTA_3KR_-FLAG er.* The same representative transgenic lines were observed. Images were taken under the same magnification. Scale bar: 25 μm. **(D)** Quantitative analysis. Stomata + meristemoid index (number of stomata and meristemoids per 100 epidermal cells) of the cotyledon abaxial epidermis from 7-day-old seedlings of respective genotypes (n = 10). One-way ANOVA followed by Tukey’s HSD test was performed, and statistically different groups are labeled with distinct letters (e.g., a, b, c, d). **(E)** Representative confocal microscopy of cotyledon abaxial epidermal from 7-day-old wild type (WT), *er-105* (*er*), *ERECTA-YFP er*, and *ERECTA_3KR_-YFP er.* The same representative transgenic lines were observed. Images were taken under the same magnification. Scale bar: 25 μm. **(F)** Quantitative analysis. Stomata + meristemoid index (number of stomata and meristemoid per 100 epidermal cells) of the cotyledon abaxial epidermis from 7-day-old seedlings of respective genotypes (n = 10). One-way ANOVA followed by Tukey’s HSD test was performed, and statistically different groups are labeled with distinct letters (e.g., a, b, c, d).

### K63-linked polyubiquitination of ERECTA by PUB30 and PUB31 is reduced in the predicted ubiquitination site mutant versions

To confirm that these lysine residues are the major ubiquitination sites, we first performed *in vitro* ubiquitination assays with recombinant ERECTA cytoplasmic domain fused with maltose binding protein (MBP-ERECTA_CD) as well as its triple lysine-to-arginine substitution version (MBP-ERECTA_CD_3KR_) as substrates. The ubiquitination of MBP-ERECTA_CD_3KR_ by PUB30/PUB31 was much weaker than that of MBP-ER_CD (Fig. 2A, S4A), suggesting that PUB30/PUB31 ubiquitinates ERECTA at the K617, K625, and K668 residues. E2 ubiquitin–conjugating enzymes (UBCs) act as key mediators of ubiquitin chain assembly and E3 ligases show pairing specificity with different E2s (Kraft *et al*., 2005; Turek *et al*., 2018; Trujillo, 2021). We then tested the auto-ubiquitination of PUB30 by UBC35, a major E2 for K63-linked polyubiquitination of plant membrane proteins (Turek *et al*., 2018; Saeed *et al*., 2023), and UBC8, which has been routinely utilized to activate PUB12/13 for ubiquitination assays of FLS2 and BRI1 (Lu *et al*., 2011; Zhou *et al*., 2018). Both UBC35 and UBC8 exhibited comparable activity for auto-ubiquitination of PUB30 (Fig. S4B), suggesting that E2 enzyme specificity may be more restricted under *in vivo* conditions. To address the *in vivo* role of these lysine residues in regulating ERECTA, we next compared the *in vivo* ubiquitination status of ERECTA and ERECTA_3KR_ in *er-105*, complemented with *ERECTApro::ERECTA-YFP* and *ERECTApro::ERECTA_3KR_-YFP* seedlings, respectively (Fig. 2B). Total proteins were extracted and subjected to anti-GFP immunoprecipitation, followed by immunoblotting with K48- and K63-linkage specific polyubiquitin antibodies, respectively. Both wild-type ERECTA-YFP and ERECTA_3KR_-YFP exhibited a similar level of K48-linked poly-ubiquitination. In contrast, ERECTA_3KR_-YFP showed a markedly reduced level of K63-linked poly-ubiquitination (Fig. 2B), which serves as a well-established signal for the internalization of membrane proteins in plants (Saeed *et al*., 2023).

**Figure 2.**
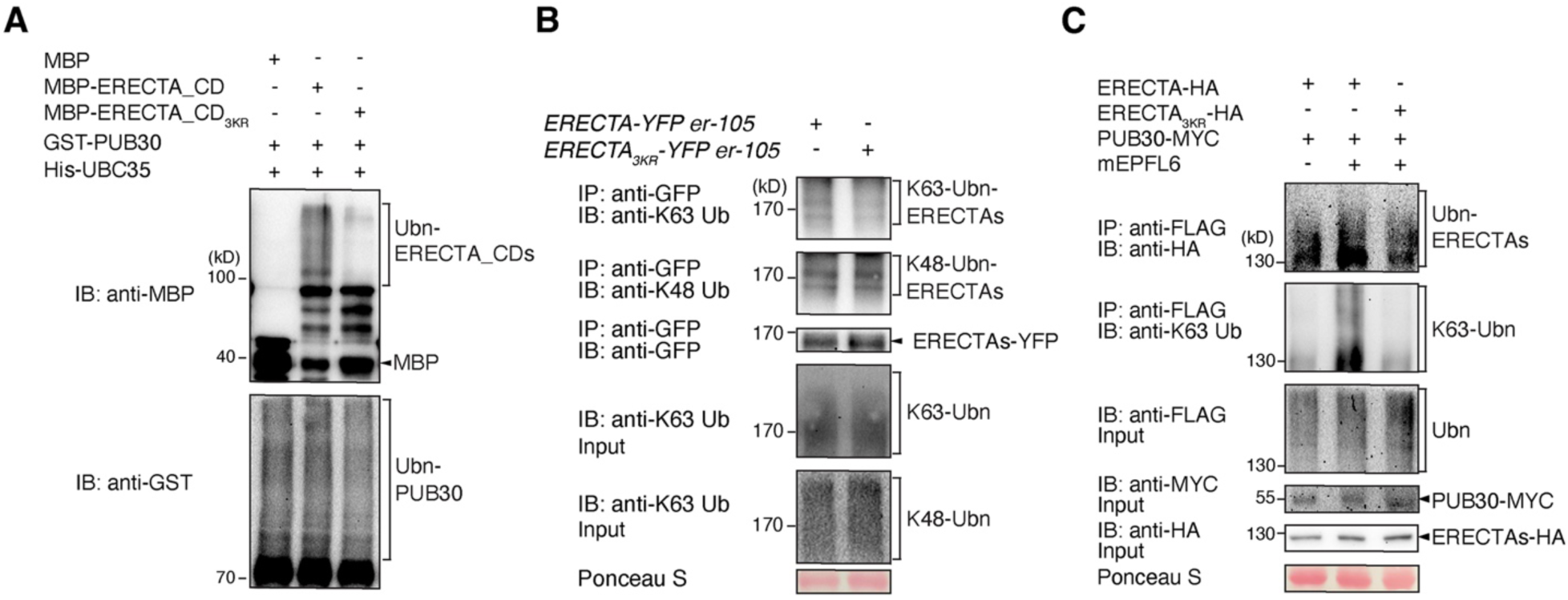
The predicted ubiquitination sites are required for the K63-linked polyubiquitination of ERECTA. (A) Lysines K617, K625, and K668 are required for the ubiquitination of ERECTA by PUB30 *in vitro*. The ubiquitination of MBP-ERECTA_CD or MBP-ERECTA_CD_3KR_ was carried out by using GST-fused PUB30 as the E3 ligase, His-fused AtUBA1 as E1 activating enzyme, and His-fused UBC35 as E2 conjugating enzyme. (B) Reduced K63-linked polyubiquitination in ERECTA_3KR_ variant. IP was performed using α-GFP antibodies on solubilized microsomal fraction protein extracts from homozygous plants expressing ERECTA-YFP or ERECTA_3KR_-YFP and the immunoblots (IB) were probed with anti-K63-linked ubiquitin and anti-GFP antibodies, respectively. (C) Lysines K617, K625, and K668 are required for the ubiquitination of ERECTA by PUB30 *in vivo*. Arabidopsis protoplasts were cotransfected with HA-tagged ERECTAs (ERECTA-HA or ERECTA_3KR_-HA), FLAG-tagged ubiquitin (FLAG-UBQ), and together with MYC-tagged PUB30 and incubated for 10 h followed by treatment with 5 μM mEPFL6 for 1 h in the presence of 5 μM MG132. After immunoprecipitation using anti-FLAG beads, the ubiquitinated ERECTA or ERECTA_3KR_ was probed with α-HA antibody. The total ubiquitinated proteins were probed by an α-FLAG antibody and PUB30 protein was probed by an α-MYC antibody. The input levels of ERECTA or ERECTA_3KR_ were probed with α-HA antibody.

To address whether this K63-ubiquitination of ERECTA is triggered by the ligand perception, we further carried out *in vivo* ubiquitination assays in *Arabidopsis* protoplasts upon bioactive mature EPFL6 (mEPFL6) peptide treatment. Laddering bands with high molecular mass proteins were observed following immunoprecipitation (Fig. 2C), indicating that ERECTA undergoes K63 polyubiquitination *in vivo*. Strikingly, the ubiquitination of ERECTA_3KR_-HA by PUB30 was significantly reduced compared with wild-type ERECTA-HA (Fig. 2C). Notably, the K63 linkage-specific ubiquitination of ERECTA_3KR_-HA by PUB30 was impaired (Fig. 2C). Based on these results, we conclude that the three lysine residues within the cytoplasmic domain of ERECTA, K617, K625, and K668, are the major K63-linked ubiquitination sites of ERECTA by its E3-ligase PUB30/PUB31 and responsible for its eventual degradation.

### Ubiquitination-deficient ERECTA displays reduced ligand-induced endocytosis

Modification of plasma membrane proteins with K63-ubiquitin chains impacts their endocytosis (Foot *et al*., 2017; Aniento *et al*., 2021; Saeed *et al*., 2023). Based on our results, we hypothesized that the K63-ubiquitination sites of ERECTA regulate its endocytosis. We therefore examined the subcellular behaviors of ERECTA-YFP as well as its ubiquitination-deficient version ERECTA_3KR_-YFP. As shown in Fig. 3A, C, we detected ERECTA-YFP-labeled punctae that co-localized with FM4-64, a styryl dye for tracing endocytic pathways in plants (Meckel *et al*., 2004), suggesting that ERECTA is actively undergoing endocytosis. Compared to the wild-type ERECTA-YFP, ERECTA_3KR_-YFP exhibit strong, brighter YFP fluorescence signals at the plasma membrane owing to its increased stability (Fig. 3A, B). Nevertheless, the number of ERECTA_3KR_-YFP endosomes is significantly reduced (Fig. 3B, C). Consistently, the signal ratio of PM to endosomes increased strikingly in ERECTA_3KR_-YFP compared with ERECTA-YFP (Fig. 3A-D).

**Figure 3.**
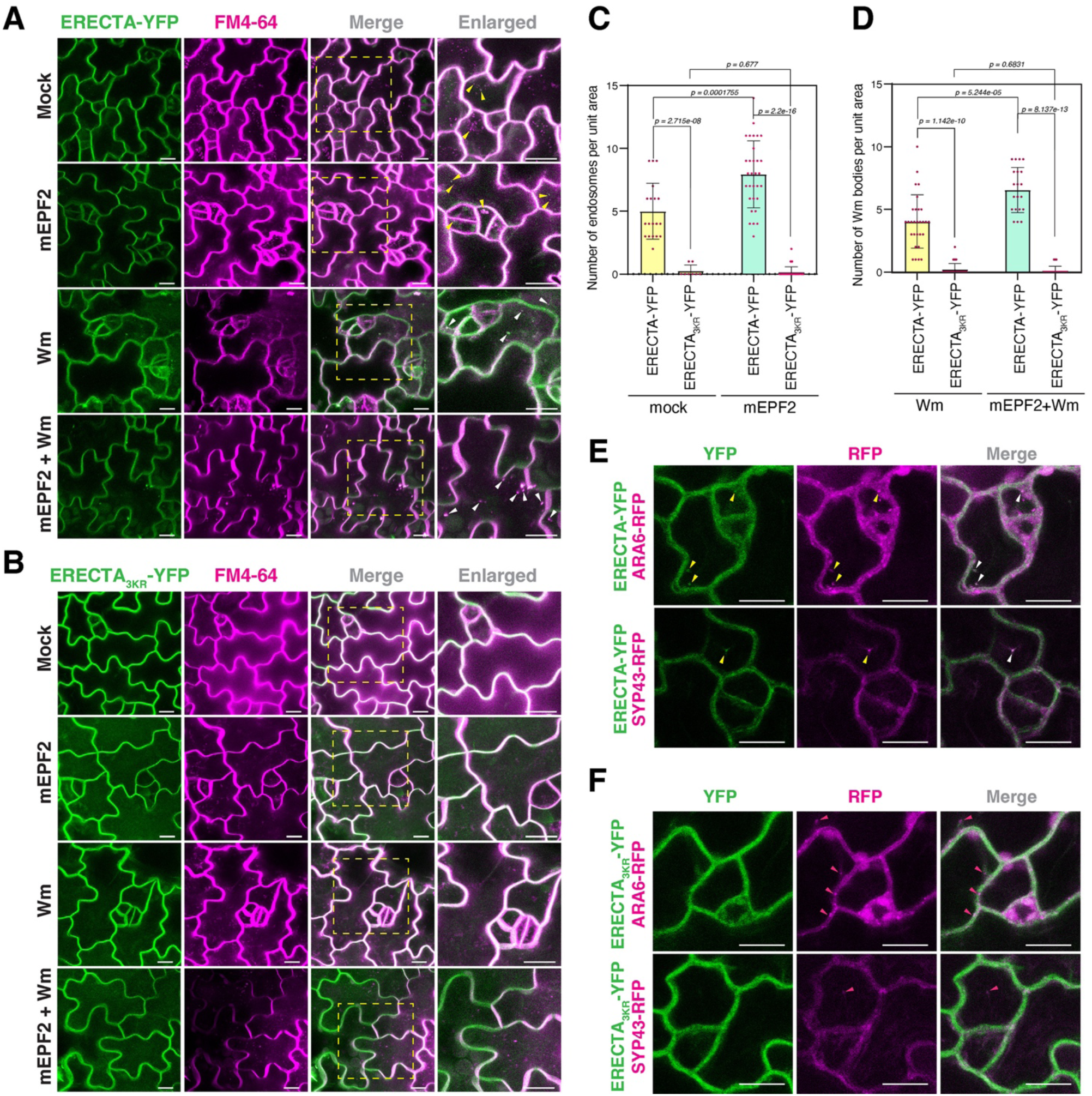
The ubiquitination of ERECTA is required for ligand-induced endocytic trafficking. **(A,B)** Representative confocal images of ERECTA-YFP (A) and ERECTA_3KR_-YFP (B) in epidermis of 4-day-old seedlings co-stained with 8 µM FM4-64 (magenta), following 5 µM mEPF2 peptide and 30 µM Wm treatment for 90 min. Arrowheads in yellow indicate ERECTA-YFP endosomes co-localizing with FM4-64 stains. Arrowheads in white indicate Wm bodies of FM4-64 co-localizing with ERECTA-YFP. The enlarged views are shown in the yellow dashed boxes. Scale bars:10 µm. **(C)** Quantification of endosome number per unit area (82 µm x 82 µm) in ERECTA-YFP and ERECTA_3KR_-YFP with or without 5 µM mEPF2 treatment. Each dot represents number of endosomes per image. Welch’s unpaired t-tests were performed, and the corresponding p-values were labeled on the plots. Number of images analyzed for mock and mEPF2, n = 18, 32, 9, 33, respectively. **(D)** Quantification of endosome number per unit area in ERECTA-YFP and ERECTA_3KR_-YFP seedlings treated with 30 µM Wm, along with or without 5 µM mEPF2 treatment. Each dot represents number of endosomes per image. Welch’s unpaired t-tests were performed, and the corresponding p-values were labeled on the plots. Number of images analyzed for Wm and Wm + mEPF2, n = 29, 36, 20, 24, respectively. **(E, F)** Representative confocal images showing co-localization of ERECTA-YFP (E) and ERECTA_3KR_-YFP (F) analysis with MVB/LE marker ARA6 (top) and TGN/EEs marker SYP43 (bottom) in the epidermis of 4-day-old seedlings. Merged images are shown in the right panel. Yellow arrows indicate the endosomes bearing ERECTA-YFP, ARA6-RFP and SYP43-RFP. White arrows point to the co-localization between ERECTA-YFP and the markers. Red arrows indicate the endosomes labeled only with the markers, without ERECTA_3KR_-YFP. Scale bars:10 µm.

To delineate which specific subcellular route is impacted by the loss of proper K63-ubiqutination of ERECTA, we next performed a pharmacological approach using Wortmannin (Wm), an inhibitor of phosphatidylinositol-3 (PI3) and phosphatidylinositol-4 (PI4) kinases (Foissner *et al*., 2016). Wm treatment causes a fusion of provacuolar compartments to form multi-vesicular bodies (MVB), also known as Wm bodies (Wang *et al*., 2009). The Wm application resulted in the formation of typical ring-like Wm bodies marked by both ERECTA-YFP and FM4-64 (Fig. 3A, D). Notably, much fewer Wm bodies were observed in ERECTA_3KR_-YFP seedlings than the WT ERECTA-YFP background (Fig. 3B, D).

To define the subcellular localization of endocytosed ERECTA-YFP and the specific location impacted by the lack of K63-ubiquitination, we next examined co-localization of ERECTA with intracellular trafficking marker proteins. Specifically, we observed co-localization of wild-type and K63-ubiquitination-deficient ERECTA-YFP fusions with RFP-ARA6 for MVB/LE (late endosomes) and plasma membrane and SYP43-RFP for trans-Golgi network/ early endosomes (TGN/EEs) (Ebine et al., 2011; Postma et al., 2016). ERECTA-YFP extensively co-localizes and moves together with RFP-ARA6 (Fig. 3E). Similarly, ERECTA-YFP-labeled punctae are also labeled by SYP43-RFP (Fig. 3F). Notably, much reduced co-localization signals were observed in ERECTA_3KR_-YFP seedlings (Fig. 3E, F), further confirming that endocytosed ERECTA-YFP predominantly resides on the TGN/EEs and MVB/LE and that K63-linked ubiquitination of ERECTA is essential for its endocytosis.

It has been reported that EPF1 and EPFL6, ligands of ERECTA-family receptors, trigger the internalization of ERL1 and that EPF2/EPFL6 application triggers the ubiquitination of ERECTA by PUB30/31 (Qi et al., 2020; Chen et al., 2023). We therefore asked whether the ligand-induced endocytosis of ERECTA depends on its ubiquitination. For this purpose, we took advantage of the biologically active mature EPF2 (mEPF2) peptides (Fig. 3A, B; Lee et al., 2012; Qi et al., 2017, 2020; Chen et al., 2023). The application of mEPF2 intensified the number of ERECTA-YFP-labeled endosomes per area, especially more evident when the ligand was applied together with Wm (Fig. 3A, C, D). In contrast, the ligand application failed to induce ERECTA_3KR_-YFP-labeled endosomes (Fig. 3B-D), further suggesting that ubiquitination of ERECTA is a prerequisite for its ligand-induced endocytosis.

### K63-linked ubiquitination is required for eventual vacuolar degradation of ERECTA

Receptors that are internalized through MVB/LE pathways are eventually targeted for degradation in the lytic vacuole (Reyes *et al*., 2011). We examined whether the ubiquitination-induced endocytosis of ERECTA is the major path for ERECTA protein degradation *in vivo*. Consistent with previous results (Fig. 3), higher accumulation of ERECTA_3KR_ (ERECTA_3KR_-YFP) was detected compared with the wild-type ERECTA (ERECTA-YFP) (Fig. 4A, B). By contrast, the transcriptional levels were not significantly different between wild-type ERECTA and ERECTA_3KR_ seedlings, suggesting that the effects of ERECTA_3KR_ accumulation are indeed posttranslational.

**Figure 4.**
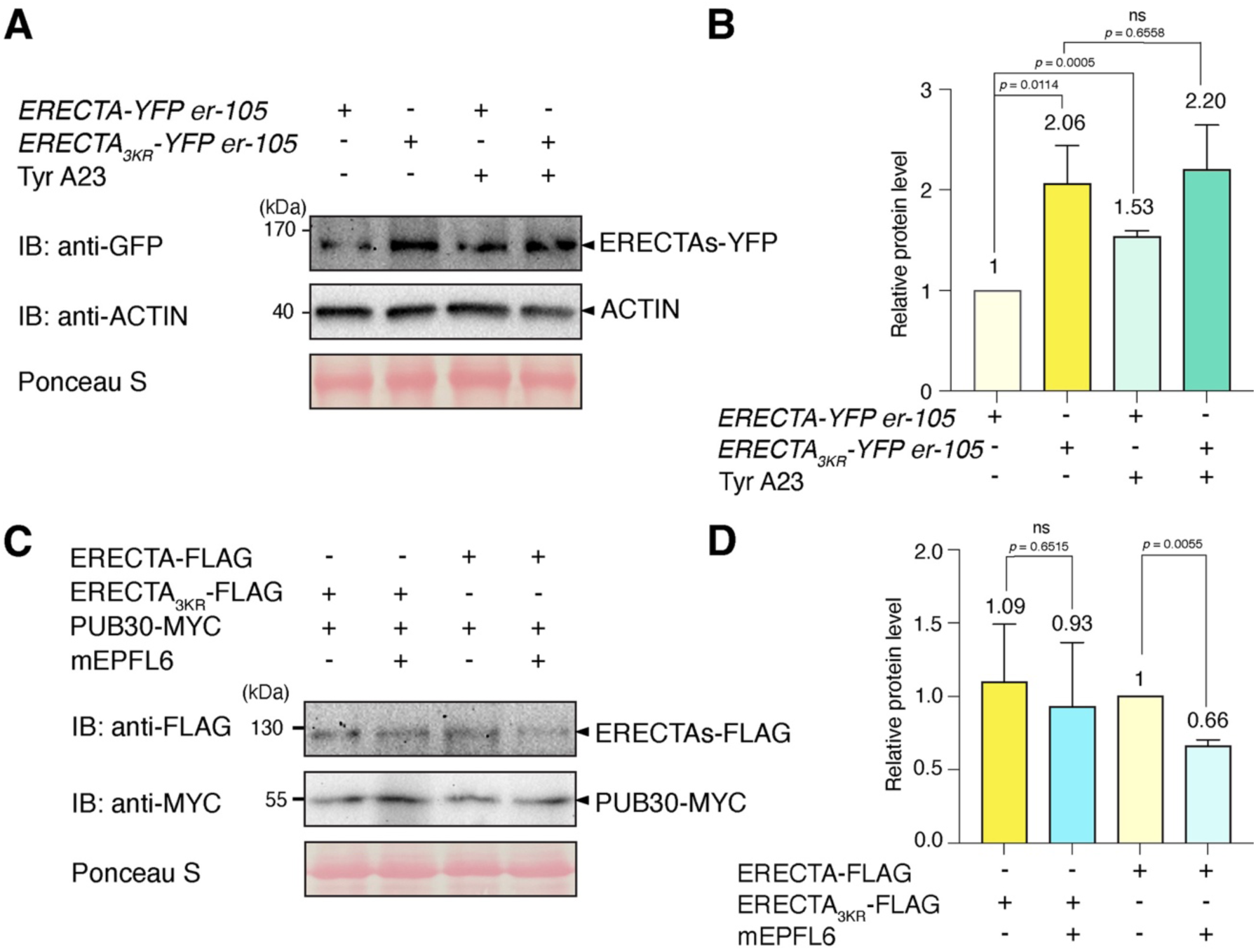
K63-linked ubiquitination is required for eventual vacuolar degradation of ERECTA. **(A)** Protein accumulation in *ERECTA-YFP er* and *ERECTA_3KR_-YFP er*, in the absence and presence of the endocytosis inhibitor Tyrphostin A23 (Tyr A23). Total proteins were isolated from 5-day-old seedlings and probed by an α-GFP antibody. The protein inputs were equilibrated using α-Actin antibodies. **(B)** Quantification of protein abundance (ERECTA/Actin and *ERECTA_3KR_*/Actin) (n = 3 biological replicates) in the absence and presence of Tyr A23. Welch’s unpaired t-tests were performed, and the corresponding p-values were labeled on the plots. **(C)** Lysines K617, K625, and K668 are required for the destabilization of ERECTA-FLAG after mEPFL6 application in Arabidopsis protoplasts co-expressing FLAG-tagged ERECTAs (ERECTA-FLAG or ERECTA_3KR_-FLAG) together with MYC-tagged PUB30. Protoplasts expressing the indicated proteins were treated with 50 μM CHX and 5 μM mEPFL6 for 3 h before total protein was examined with immunoblot. **(D)** Quantification of protein abundance (ERECTA/Rubisco and *ERECTA_3KR_*/Rubisco) (n = 3 biological replicates) in the absence and presence of mEPFL6. Welch’s unpaired t-tests were performed, and the corresponding p-values were labeled on the plots.

To further assess whether endocytic/vacuolar degradation pathways are responsible for the enhanced protein abundance of ERECTA_3KR_-YFP, we subsequently treated the seedlings with Tyrphostin A23 (Tyr A23), an inhibitor of clathrin-mediated endocytosis (Santuari *et al*., 2011) and Concanamycin A (Con A), a vacuolar ATPase inhibitor that is known to diminish protein degradation in the lytic vacuole (Kleine-Vehn *et al*., 2008). As shown in Fig. 4A, B, and Fig. S5, treatments of both Tyr A23 and Con A resulted in a significant increase in ERECTA protein accumulation, but not in that of the K63-ubiquitination deficient ERECTA_3KR_. Finally, we tested the function of K63-ubiquitination on the ligand-induced degradation of ERECTA protein. For this purpose, we co-expressed ERECTA or ERECTA_3KR_, and PUB30 in Arabidopsis protoplast and performed co-treatment with mEPFL6 and cycloheximide (CHX, *de novo* protein synthesis inhibitor). Upon mEPFL6 treatment, ERECTA-FLAG showed a distinct decrease in protein level (Fig. 4C, D). In contrast, PUB30-MYC conferred a smaller reduction of ERECTA_3KR_-FLAG protein level (Fig. 4C, D). Taken together, our results thus demonstrate that ubiquitination-induced endocytosis of ERECTA triggers the degradation of the receptor.

## Discussion

Our previous study identified two PUB-type E3 ligases, PUB30 and PUB31, as key attenuators of the ERECTA regulatory circuit, which ensure optimal signal outputs. Recently, K63-ubiquitin chains have been reported to serve as a general signal for early endocytosis of integral plasma mambrane proteins, including FLS2, BRI1, and PIN1 (Saeed *et al*., 2023). However, the type of ubiquitin linkage on ERECTA and its connection to receptor turnover has remained unclear. In this study, we identify K63-linked ubiquitination of ERECTA as a key regulatory mechanism that controls its internalization and vacuolar degradation during stomatal development, as well as during inflorescence and pedicel elongation. Perception of EPF/EPFL peptides initiates a defined signaling cascade through heterodimerization with BAK1/SERKs, leading to suppression of a transcription factor that drives stomatal lineage initiation (Lampard *et al*., 2008; Horst *et al*., 2015; Meng *et al*., 2015). To preserve signal fidelity and spatial restriction, receptor activity must be tightly regulated. Our data show that PUB30/PUB31-mediated K63-linked ubiquitination promotes internalization and degradation of ERECTA upon ligand activation (Fig. 5). By mapping and validating three lysine residues essential for this modification, we show that K63-linked ubiquitination is required for ligand-induced endocytosis and subsequent protein degradation of ERECTA, thereby preventing excessive or prolonged receptor activity. These findings offer mechanistic insight into how post-translational control of plasma membrane-localized RLKs contributes to precise signaling during plant development.

**Figure 5.**
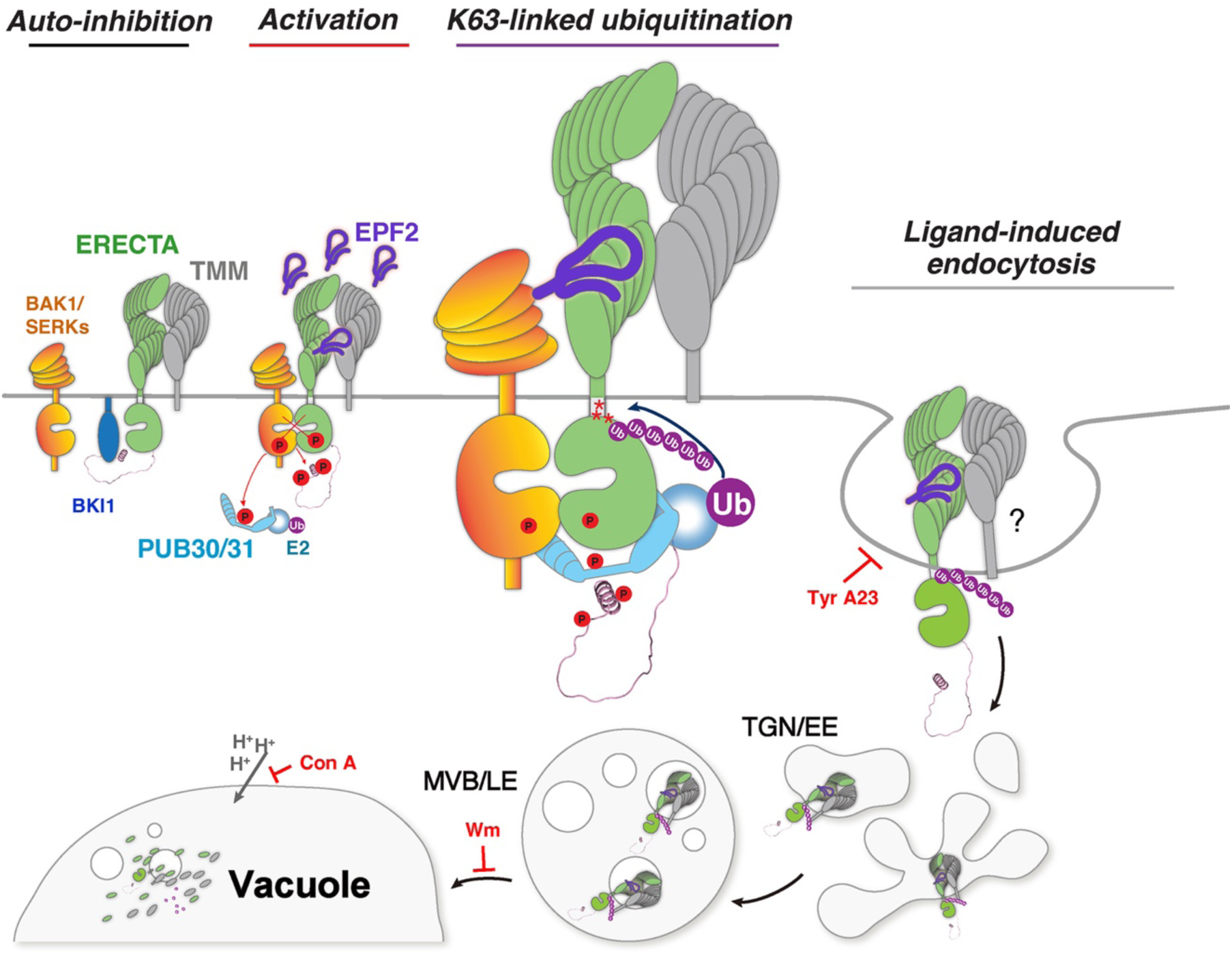
Schematic model of ERECTA endocytosis and homeostasis regulated by ligand-induced ubiquitination during stomatal development. Initially, BKI1 (blue) associates with ERECTA (green) in the absence of ligand, and ERECTA is in the basal state (*Auto-inhibition*). Upon perception of the ligand EPF2 (violet), ERECTA signaling becomes activated. The formation of the receptor complex, ERECTA and its co-receptors TMM (gray) and BAK1/SERKs (orange), initiates signal transduction that ultimately restricts stomatal development (*Activation*). Following activation, ERECTA is ubiquitinated at lysine residues K617, K625, and K668 via PUB30/31 (cyan) (*K63-linked polyubiquitination*). Thereafter, the ubiquitinated ERECTA undergoes endocytosis and vacuolar/lysosomal degradation, which maintains appropriate receptor homeostasis to ensure proper stomatal development (*Ligand-induced endocytosis*). Whether TMM (gray) co-internalizes with ERECTA upon ligand perception remains to be investigated.

Our sequence alignment revealed that the location of BRI1 ubiquitination site (K668) within its juxtamembrane domain is not conserved in ERECTA (Fig. 1A). Nevertheless, our site-directed mutagenesis of the lysine residues within the juxtamembrane of ERECTA (K617 and K625) together with the known ubiquitination site conferred the stable protein accumulation and hyperactivity of ERECTA signaling. These results support a role for the juxtamembrane domain of ERECTA as a ubiquitination targeted domain in controlling receptor stability. Consistently, we previously observed that the truncated ERECTA protein lacking the entire cytoplasmic domain, including the juxtamembrane domain, stably accumulates at the plasma membrane and confers strong dominant-negative effects (Shpak *et al*., 2003). Likewise, the dominant-negative ERL1, which also lacks the entire cytoplasmic domain, triggers a stable accumulation at the plasma membrane (Qi *et al*., 2020), suggesting that the ubiquitination-dependent regulation occurs in other ERECTA-family RKs. Further structural analyses may elucidate the modular functions of ERECTA cytoplasmic domains (Fig. 1B).

The ubiquitin-mediated trafficking of ERECTA bears mechanistic similarity to that of BRI1 and FLS2, which are also subject to K63-linked polyubiquitination by the distinct members of the U-box E3-ligases PUB12/PUB13 (Zhou *et al*., 2018; Saeed *et al*., 2023). It is highly likely that these PUB-type E3 ligases pair up with E2 ubiquitin-conjugating enzymes (e.g., UBC35/UBC36) to mediate K63-linked ubiquitination (Saeed *et al*., 2023), endocytosis, and subsequent receptor degradation, thereby pointing to a shared regulatory framework across RLKs. However, the developmental role of ERECTA imposes stricter spatial constraints. ERECTA interprets multiple EPF/EPFL peptides to control stomatal patterning, a process that requires signaling to be confined both spatially and temporally, often at the level of individual cell files. Our data show that PUB30/PUB31-mediated attenuation of ERECTA via ubiquitination, receptor internalization and vacuolar degradation, is essential for maintaining positional fidelity and preventing ectopic activation (Figs. 3, 4). In contrast, FLS2 responds to broadly diffusible pathogen-associated signals to trigger acute immune responses (Chinchilla *et al*., 2007). Although BRI1 signaling shows tissue-specific outputs depending on developmental context (Planas-Riverola *et al*., 2019; Blanco-Touriñán *et al*., 2024), its endocytosis has been studied using seedling roots, where BRI1 is uniformly expressed (Di Rubbo *et al*., 2013; Zhou *et al*., 2018; Saeed *et al*., 2023). It has become evident that tissue- and cell-type-specific regulation of signaling events are critical for immunity and development (Sparks *et al*., 2013; Fukuda & Hardtke, 2019; Han & Tsuda, 2023). As such, the knowledge obtained through this study will help unravel how receptor trafficking defines positional information in plants.

There has been a debate over whether plant receptors signal exclusively at the plasma membrane or if their signaling activity is sustained in endosomes after the ligand-induced internalization, similar to what is observed for the EPIDERMAL GROWTH FACTOR (EGF) receptor kinases in mammals (Lai *et al*., 1989; Pinilla-Macua *et al*., 2025). In the case of BRI1, early evidence suggested its endosomal signaling (Geldner *et al*., 2007). A subsequent study presented that increasing BRI1 population in the TGN/EE did not affect BR signaling, thus suggesting that BRI1 signals from the plasma membrane (Irani *et al*., 2012; Zhang *et al*., 2019). Our study shows that the K63-ubiquitination-deficient mutant of ERECTA (ERECTA_3KR_) stably accumulates at the plasma membrane and severely compromises ligand-induced endocytosis (Fig. 3). Importantly, this severe reduction of ERECTA endocytosis results in hyperactivity of ERECTA signaling, leading to over-inhibition of stomatal development as well as excessive elongation of pedicels (Figs 1, S1-2). Taken together, our work provides evidence supporting that ERECTA signals at the plasma membrane.

Several key questions remain unresolved. How ERECTA trafficking interfaces with broader endomembrane processes (Banjade *et al*., 2019), as well as how distinct EPF/EPFL peptide ligands and co-receptors (e.g. TMM, BAK1/SERKS: Fig. 5) influence the dynamics of ERECTA endocytosis, remains to be clarified. Likewise, how ERECTA ubiquitination cooperates with or antagonizes other post-translational modifications, such as phosphorylation, to form combinatorial regulatory outputs is an important area for future investigation (Zhang & Zeng, 2020). Moreover, it has not been established whether environmental or hormonal cues modulate ERECTA ubiquitination, thereby linking external signals to spatial control of stomatal development (Qi & Torii, 2018). Finally, the K63-linked ubiquitination attenuates multiple receptor kinases, including ERECTA, BRI1, and FLS2, pointing to a conserved mechanism that limits receptor longevity in contexts requiring tight signaling control (Saeed *et al*., 2023). Comparative analyses across species and RLK subclasses may uncover shared principles underlying membrane receptor regulation in plants. Together, these questions point toward a broader framework for understanding how signaling precision is achieved through layered post-translational control of receptor homeostasis.

## Materials and Methods

### Plant materials and growth conditions

The Arabidopsis accession Columbia (Col) was used as the wild type. All plants used in this study are in a Col background. The following mutants and transgenic plant lines were previously reported: *er-105*; *pub30pub31*; *ERECTApro::gERECTA-FLAG* in *er-105 (Chen et al., 2025)*. Arabidopsis seeds were surface sterilized with 30% bleach and grown on half-strength Murashige and Skoog media containing 1× Gamborg Vitamin (Sigma), 0.75% Bacto Agar and 1% w/v sucrose for 9 days and then transplanted into soil. Plants were grown under long-day conditions (16 h light/8 h dark) at 22 °C.

### Plasmid construction and generation of transgenic plants

For recombinant protein expression, the following plasmids were previously reported: pJA51 (MBP-ERECTA_CD), pCLL107 (GST-PUB30), and pCLL109 (GST-PUB31) (Chen *et al*., 2023; Chen *et al*., 2025). The following plasmids were generated: pCLL285 (MBP-ERECTA_CD_K617RK625RK668R_, MBP-ERECTAR_CD_3KR_). Site-directed mutagenesis was performed using a 2-sided PCR overlap extension followed by assembly into linearized vector pMALC2. For complementation assays, the following plasmids were generated: pCLL284 (*ERECTApro::gERECTA_K668R_-FLAG*), pCLL200 (*ERECTApro::gERECTA_K625RK668R_-FLAG*), pCLL201 (*ERECTApro::gERECTA_K617RK668R_-FLAG*), pCLL287 (*ERECTApro::gERECTA_K617RK625RK668R_-FLAG*), pCLL283 (*ERECTApro::gERECTA-YFP*), pCLL288 (*ERECTApro::gERECTA_K617RK625RK668R_-YFP*). A three-way Gateway system was utilized to generate a series of ERECTA constructs driven by the ERECTA promoter. See Tables S1 and S2 for details of plasmid and oligo DNA information. Plasmids were transformed into Agrobacterium GV3101/pMP90 and subsequently into Arabidopsis by floral dipping. Over 10 lines were characterized for the phenotypes and reporter gene expressions.

### Microscopy for stomatal phenotype

Confocal microscope images for stomatal phenotype were taken as previously described using Leica SP5-WLL operated by LAS AF (Leica). Cell peripheries of seedlings were visualized with propidium iodide (Molecular Probes). Fluorescence signal was detected under the following conditions: Propidium iodide, excitation at 561 nm and the emission range from 582 to 617 nm. For internalization imaging ERECTA-YFP, ERECTA_3KR_-YFP and other membrane organelle markers, Z-stack images were acquired at 0.49 µm intervals to cover the entire epidermal cell depth at a resolution of 1024 x 1024 pixels. Maximum intensity projections were generated using Leica LAS AF software and analyzed with Fiji (https://imagej.net). Images were captured using either a 20x water-immersion objective for stomatal observation or a 63x water-immersion objective for endocytosis assays. Fluorescence signals were detected under the following conditions: YFP, excitation 514 nm and emission range from 518 to 550 nm; RFP and FM4-64, excitation 555 nm and emission range from 573 to 630 nm. For endosome-labeled FM4-64 quantification, the maximum projection of Z-stack images was then generated by Leica LAS FA software. For Wortmannin treatment, seedlings were incubated with 30 µm Wortmannin (in 0.25 % DMSO) for 90 min with gentle shaking. A mock control consisted of 0.25% DMSO in half-strength MS liquid. For mEPF2 treatment, the purified peptide solution was diluted to 5 µM using half-strength MS liquid and incubated with seedlings for 90 min under gentle shaking, followed by staining with 8 µM of FM4-64. Endosomes were defined as structures with diameters of 0.2-0.5 µm, while Wm bodies were defined as punctae larger than 0.5 µm. Each experiment was repeated at least three times, each with at least three seedlings per repeat. The confocal images were false colored, and brightness/contrast were uniformly adjusted using Photoshop 2022 (Adobe).

### Quantitative analysis and statistics

For analysis of the epidermis, abaxial cotyledons from 7-day-old seedlings of relevant genotypes were subjected to PI staining and confocal microscopy. The central regions overlying the distal vascular loop were imaged, and the number of epidermal cells, stomata, and meristemoids was quantified. Pedicel lengths were measured using ImageJ. Statistical analysis was performed using R ver. 4.4.0 operated under R-Studio ver. 2025.05.0+496 (https://www.rstudio.com), and graphs were generated using R ggplot2 package. For all violin plots, the shape of each violin represents the kernel density estimation of the data distribution across its range, with the width at each vertical position corresponding to the local data density. Each dot represents a jittered individual data point. For genotype-phenotype comparison, One-way ANOVA followed by Tukey’s HSD test was performed for comparisons among multiple groups. Statistical differences were indicated by different letters (e.g., a, b, c, d). For individual sample size (n), see corresponding figures or figure legends.

Gene expression levels in RT-qPCR analysis, quantification of endosomes, and protein abundance from Immunoblot assays were analyzed using GraphPad Prism software (version 10.3.0 (461), GraphPad Software). For comparisons between matched samples, Welch’s unpaired t-tests were performed. For individual sample size (n) and p values, see corresponding figures or figure legends.

### Expression, purification, and refolding of peptides

Recombinant mEPFL6 peptide was prepared as described previously (Lee *et al*., 2012). Bioactivities of the refolded mEPFL6 peptide were confirmed as described previously (Lee *et al*., 2012).

### *In vitro* and *in vivo* ubiquitination assays

The *in vitro* ubiquitination reactions contained 1 μg each of substrate (MBP-ERECTA_CD or MBP-ERECTA_CD_3KR_), HIS-E1 (AtUBA1), HIS-E2 (AtUBC8 or AtUBC35), HIS-ubiquitin (Boston Biochem) and GST-PUB30 in the ubiquitination reaction buffer (0.1 M Tris–HCl, pH 7.5, 25 mM MgCl_2_, 2.5 mM dithiothreitol and 10 mM ATP; the final volume 30 μL). The reactions were incubated at 30 °C for 3 h, stopped by adding 4x SDS sample loading buffer and boiled at 95 °C for 5 min. The samples were then separated by 8% SDS–PAGE, and the ubiquitinated ER_CDs were detected by immunoblot analysis with anti-MBP (E8032, 1:10,000; New England Biolabs) as the primary antibody, whereas the auto-ubiquitination was detected by immunoblot analysis with anti-GST (ab92, 1:5,000; Abcam) as the primary antibody. As a secondary antibody, goat anti-mouse IgG H&L (HRP) (ab205719, Abcam) was used at a dilution of 1:5,000. The protein blots were visualized using a Chemi-luminescence assay kit (34095, Thermo Scientific) and the Image Lab software (version 6.0.1, Bio-Rad).

For *in vivo* ubiquitination assays, fresh Arabidopsis protoplasts were co-transfected with FLAG-tagged ubiquitin (FLAG-UBQ), HA-tagged ERECTA variants and together with a control vector or MYC-tagged PUB30 and incubated for 9 h followed by treatment with 5 μM mEPFL6 for 1 h in the presence of protease inhibitor MG132 (5 μM; M7449, Sigma). The ubiquitinated ERECTA variants were detected with α-HA (ab18181, 1:1,000, Abcam), and anti-Ubiquitin (linkage-specific K63) antibody (ab179434, 1:3,000, Abcam), IB after IP with α-FLAG (ab205606, Abcam) antibody. The total proteins were detected by α-HA (ab18181, 1:1,000, Abcam), anti-FLAG (F-3165, 1:5,000, Sigma) and α-MYC (ab32, 1:1,000, Abcam) as primary antibodies. As a secondary antibody, goat anti-mouse IgG H&L (HRP) (ab205719, Abcam) was used for HA, FLAG, and MYC detection at a dilution of 1:5,000, and goat anti-rabbit IgG H&L (HRP) (ab205718, Abcam) was used for K63 ubiquitination detection at a dilution of 1:5,000. Protein blots were visualized as described in the previous section.

For *in vivo* ubiquitination level assays, the *ERECTApro::gERECTA-YFP er-105* and *ERECTApro::gERECTA_3KR_-YFP er-105* seedlings were grown vertically at 22 °C on half-strength Murashige and Skoog medium for 5 days. The immunoprecipitated (IP) was performed using α-GFP antibodies on total protein extracts from homozygous plants. The amount of IP sample loaded was normalized based on the GFP signal detected by anti-GFP immunoblotting. As secondary antibody, goat anti-mouse IgG H&L (HRP) (ab205719, Abcam) was used at a dilution of 1:5,000. Ubiquitination levels were subsequently assessed by immunoblotting with anti-Ubiquitin (linkage-specific K63) antibody (ab179434, 1:3,000, Abcam) and anti-Ubiquitin (linkage-specific K48) antibody (ab140601, 1:3,000, Abcam), respectively. As a secondary antibody, goat anti-rabbit IgG H&L (HRP) (ab205718, Abcam) was used at a dilution of 1:5,000. Protein blots were visualized as described in the previous section.

### Inhibitor treatment and immunoblot assays

The *ERECTApro::gERECTA-YFP er-105* and *ERECTApro::gERECTA_3KR_-YFP er-105* seedlings were grown vertically at 22 °C on half-strength Murashige and Skoog medium for 5 days. Thereafter, seedlings were incubated with or without endocytosis inhibitor, 50 μM Tyrphostin A23 (Tyr A23) (T7165, Sigma) for 30 min. For the vacuolar ATPase inhibitor treatment, seedlings were incubated with or without vacuolar ATPase inhibitor, 1 μM Con A (ab144227, Abcam) for 60 min. Total protein extracts were separated on an 8% SDS–polyacrylamide gel and detected by immunoblot analysis with anti-FLAG (F-3165, 1:5,000, Sigma) and anti-actin (ab230169, 1:2,000, Abcam) as primary antibodies. As a secondary antibody, goat anti-mouse IgG H&L (HRP) (ab205719, Abcam) was used at a dilution of 1:5,000. Protein blots were visualized as described in the previous section.

### Protein stability assay in protoplasts

To assess ERECTA protein stability upon ligand treatment, protoplasts co-transfected with PUB30-MYC, the wild-type ERECTA-FLAG or the ubiquitination-mutant version ERECTA_3KR_-FLAG were treated with 50 μM cycloheximide (CHX, C4859, Sigma) in the presence or absence of 5 μM mEPFL6 for 3 h. Total proteins were separated on 8% SDS-PAGE gels and transferred to the PVDF membrane (Millipore) for immunoblot analysis. ERECTA variants protein and the input PUB30 protein were detected with α-FLAG antibody (F-3165, 1:5,000, Sigma) and α-MYC antibody (ab32, 1:1,000, Abcam) as primary antibodies, respectively. As a secondary antibody, goat anti-mouse IgG H&L (HRP) (ab205719, Abcam) was used at a dilution of 1:5,000. Protein blots were visualized as described in the previous section.

### RT-qPCR analysis

Tissues for qPCR analysis were harvested at 4-day-old for established lines. RNA extraction, cDNA synthesis, and RT-qPCR were performed as described previously (Pillitteri *et al*., 2011). Transcript levels were normalized against the housekeeping gene *ACTIN* (*ACT2*). For primer DNA sequences used for RT-qPCR analysis, see Table S2.

## Acknowledgements

We thank Prof. Takashi Ueda and Prof. Kazuo Ebine for a gift of ARA6-RFP and SYP43-RFP marker lines and insightful suggestions and discussion about receptor endocytosis. We also thank the Torii Lab members, especially Dr. Krishna Sepuru and Dr. Pengfei Bai, for the discussion. This work was supported by funding from the Howard Hughes Medical Institute and the start-up fund from The University of Texas at Austin, as well as the Johnson & Johnson Centennial Chair awarded to K.U.T.

## Author Contributions

Conceptualization, L.L.C., K.U.T.; Experimental Design, L.L.C., M.V.H., K.U.T.; Performance of experiments, L.L.C., M.H.V., A.M.C., C.F.Y., K.U.T. (Biochemistry experiments, L.L.C.; Confocal microscopy, L.L.C. and M.H.V.); Data analysis and Visualization, L.L.C., M.V.H., K.U.T.; Writing-Original Draft, L.L.C. and K.U.T.; Writing-Comments and Edits, L.L.C., M.H.V., A.M.C., C.F.Y., K.U.T.; Project Administration, K.U.T.; Funding Acquisition, K.U.T.

---

**Fig. S1.**
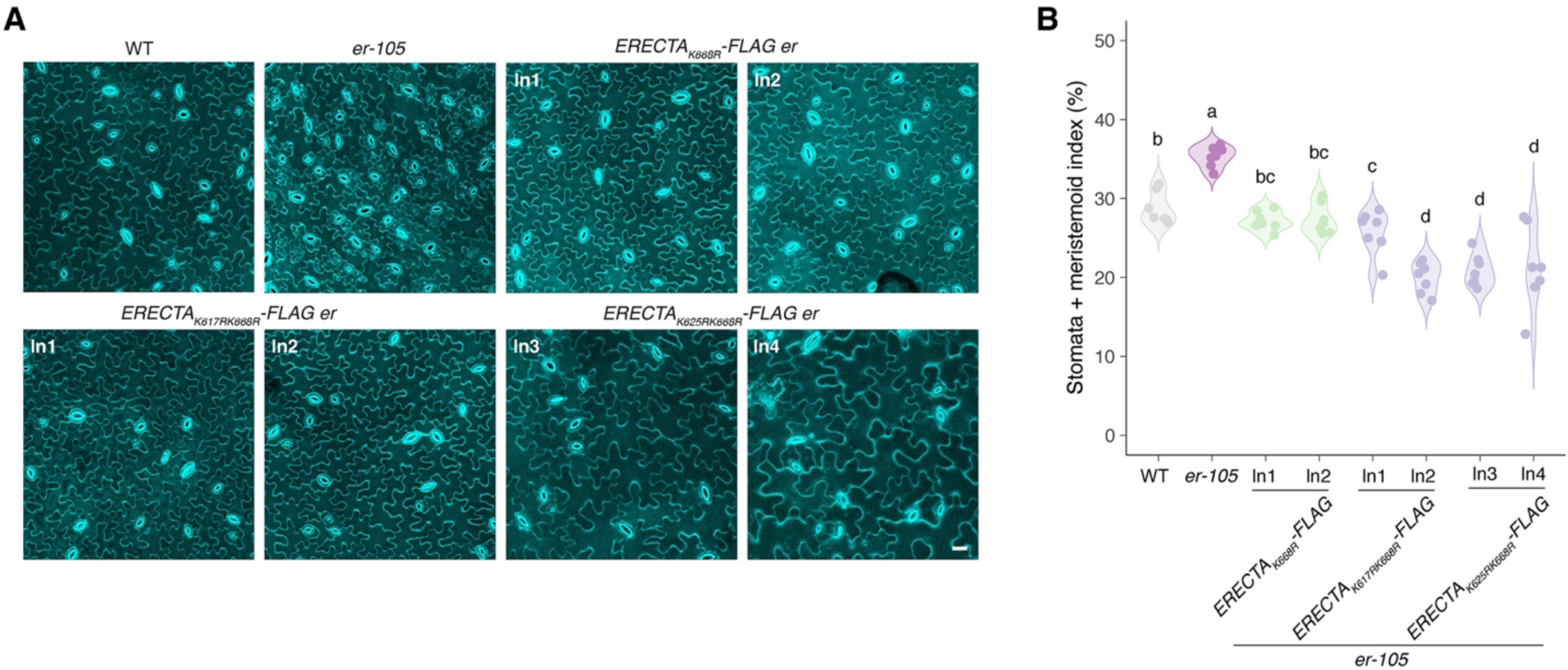
Mutations of predicted lysine residues in ERECTA show redundancy in inhibiting stomatal development, related to Fig. 1. **(A)** Representative confocal microscopy of cotyledon abaxial epidermal from 7-day-old wild type (WT), *er-105* (*er*), *ERECTA_K668R_-FLAG er*, *ERECTA_K617RK668R_-FLAG er*, and *ERECTA_K625RK668R_-FLAG er.* The same representative transgenic lines were observed. Images were taken under the same magnification. Scale bar: 25 μm. **(B)** Quantitative analysis. Stomata + meristemoid index (number of stomata and meristemoid per 100 epidermal cells) of the cotyledon abaxial epidermis from 7-day-old seedlings of respective genotypes (n = 7). One-way ANOVA followed by Tukey’s HSD test was performed, and statistically different groups are labeled with distinct letters (e.g., a, b, c, d).

**Fig. S2.**
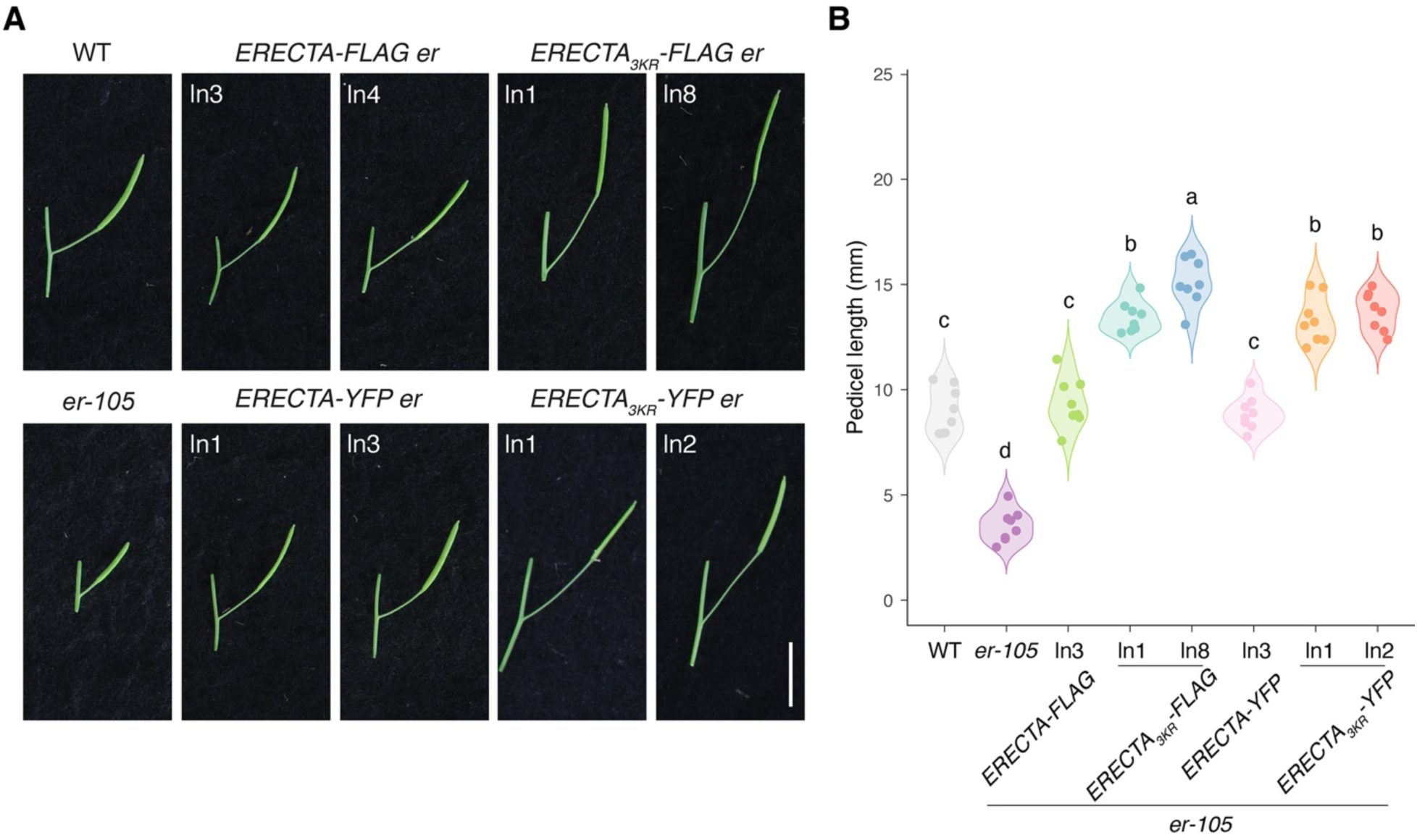
Ubiquitination-deficient ERECTA overly promotes pedicel growth, related to Fig. 1. **(A)** Representative pedicels and mature siliques of WT, *er-105* (*er*), *ERECTA-FLAG er*, *ERECTA_3KR_-FLAG er*, *ERECTA-YFP er*, *ERECTA_3KR_-YFP er* plants. For each transgenic construct, two representative lines were subjected to analysis. Images were taken under the same magnification. (Scale bar, 10 mm.) **(B)** Morphometric analysis of pedicel length from each genotype. 6-wk-old mature pedicels (n = 8) were measured. One-way ANOVA followed by Tukey’s HSD test was performed, and statistically different groups are labeled with distinct letters (e.g., a, b, c, d).

**Fig. S3.**
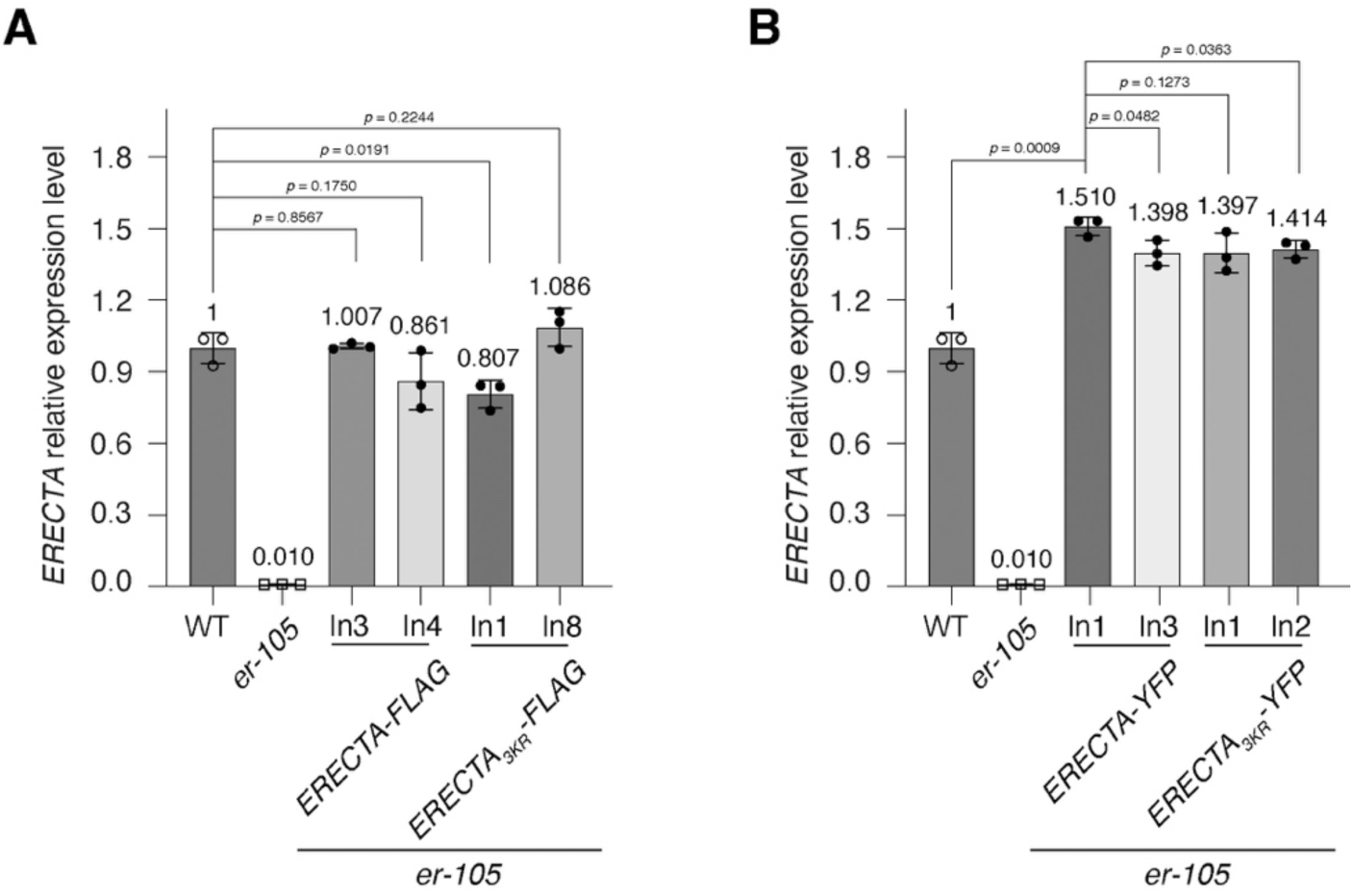
Expression level of ERECTA in ERECTA-FLAG er, ERECTA_3KR_-FLAG er, ERECTA-YFP er, ERECTA_3KR_-YFP er, related to Fig. 1. **(A)** RT-qPCR analysis of *ERECTA* in WT, *er-105* (*er*), *ERECTA-FLAG er*, *ERECTA_3KR_-FLAG er.* Transcript levels were normalized against *ACTIN* (*ACT2*) and adjusted to 1 for wild type. Bars, mean values of three technical replicates. Welch’s unpaired t-tests were performed, and the corresponding p-values were labeled on the plots. **(B)** RT-qPCR analysis of *ERECTA* in WT, *er-105* (*er*), *ERECTA-YFP er*, *ERECTA_3KR_-YFP er.* Transcript levels were normalized against *ACT2* and adjusted to 1 for wild type. Bars, mean values of three technical replicates. Welch’s unpaired t-tests were performed, and the corresponding p-values were labeled on the plots.

**Fig. S4.**
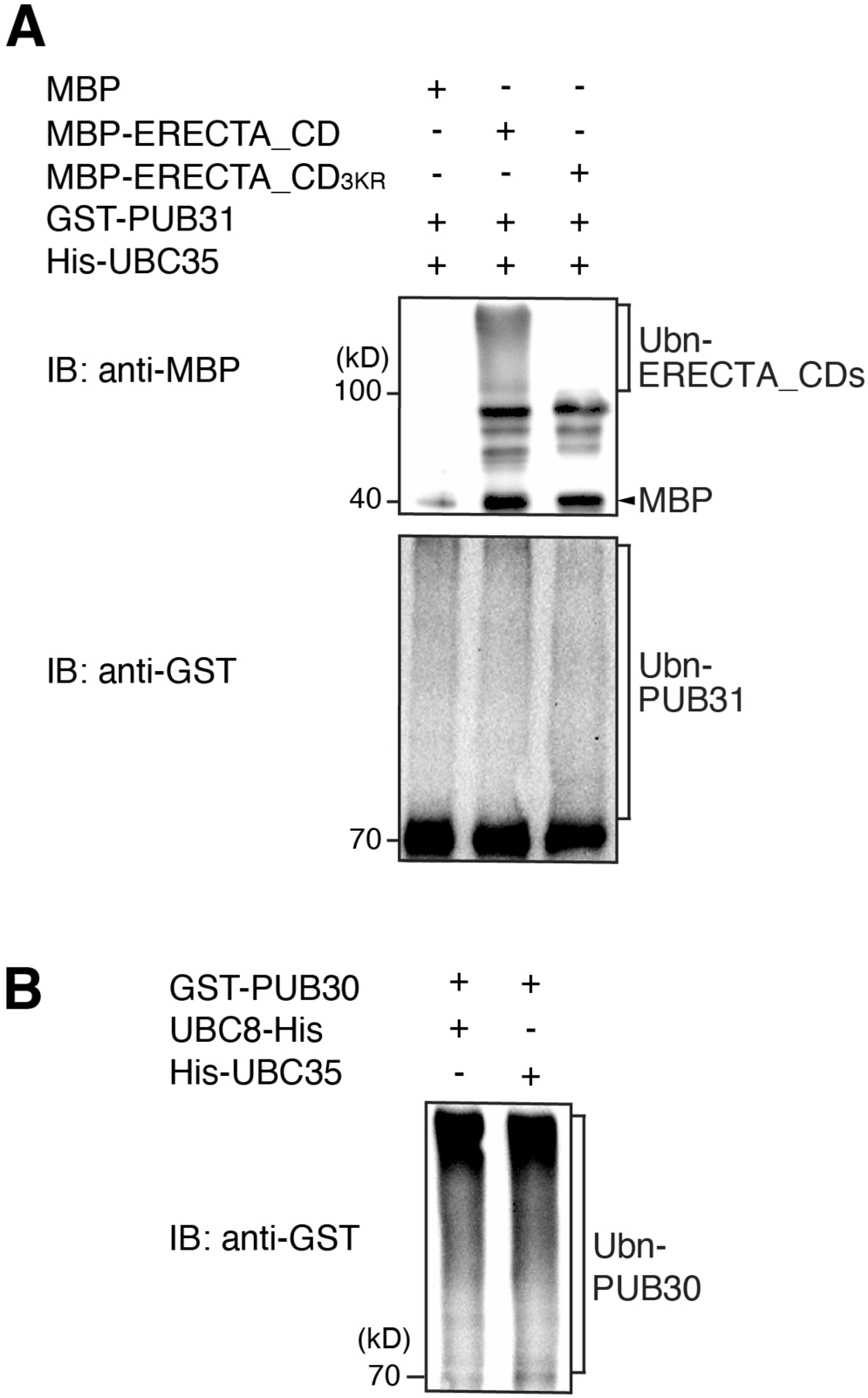
The predicted ubiquitination sites are required for polyubiquitination of ERECTA by PUB31, related to Fig. 2. **(A)** Lysines K617, K625, and K668 are required for the ubiquitination of ERECTA by PUB31 *in vitro*. The ubiquitination of MBP-ERECTA_CD or MBP-ERECTA_CD_3KR_ was carried out by using GST-fused PUB31 as the E3 ligase, His-fused AtUBA1 as E1 activating enzyme, and His-fused UBC35 as E2 conjugating enzyme. **(B)** UBC8 and UBC35 exhibit comparable activity for autophosphorylation of PUB30. The auto-ubiquitination of GST-PUB30 was performed using His-fused AtUBA1 as E1 activating enzyme, and His-fused UBC8 or UBC35 as E2 conjugating enzyme.

**Fig. S5.**
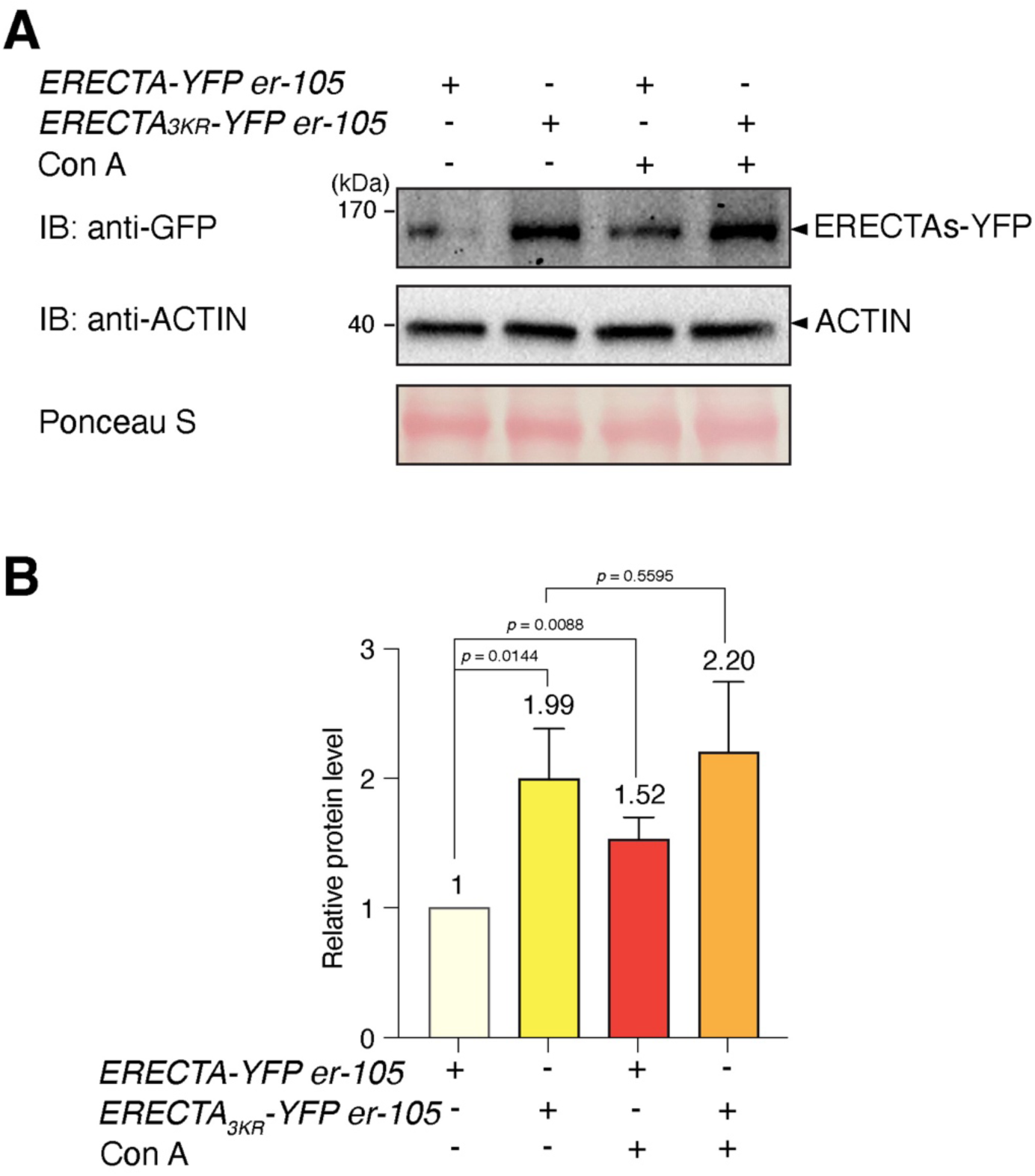
K63-linked ubiquitination is required for eventual vacuolar degradation of ERECTA, related to Fig. 4. **(A)** Protein accumulation in *ERECTA-YFP er* and *ERECTA_3KR_-YFP er*, in the absence and presence of the vacuolar ATPase inhibitor Concanamycin A (Con A). Total proteins were isolated from 5-day-old seedlings and probed by an α-GFP antibody. The protein inputs were equilibrated using α-Actin antibodies. **(B)** Quantification of protein abundance (ERECTA/Actin and *ERECTA_3KR_*/Actin) (n = 3 biological replicates) in the absence and presence of Con A. Welch’s unpaired t-tests were performed, and the corresponding p-values were labeled on the plots.

